# Data-driven Modeling of Thrombus Size and Shape in Aortic Dissections: Role of Hemodynamics

**DOI:** 10.1101/187914

**Authors:** Alireza Yazdani, He Li, Matthew R. Bersi, Paolo Di Achille, Joseph Insley, Jay D. Humphrey, George Em Karniadakis

**Affiliations:** Division of Applied Mathematics, Brown University, Providence, RI 02912; Department of Biomedical Engineering, Yale University, New Haven, CT 06520; Argonne National Laboratory, Argonne, IL 60439; Northern Illinois University, DeKalb, IL 60115

## Abstract

Aortic dissection is a pathology that manifests due to micro-structural defects in the aortic wall. Blood enters the damaged wall through an intimal tear, thereby creating a so-called *false lumen* and exposing the blood to thrombogenic intramural constituents such as collagen. The natural history of this acute vascular injury thus depends, in part, on thrombus formation, maturation, and possible healing within the false lumen. A key question is: Why do some false lumens thrombose completely while other thrombose partially or little at all? An ability to predict the location and extent of thrombus in subjects with dissection could contribute significantly to clinical decision-making, including interventional design. We develop, for the first time, a data-driven *particle-continuum* model for thrombus formation in a murine model of aortic dissection. In the proposed model, we simulate a final-value problem in lieu of the original initial-value problem with significantly fewer particles that may grow in size upon activation, representing the local concentration of blood-borne species. Numerical results confirm that geometry and local hemodynamics play significant roles in the acute progression of thrombus. Despite geometrical differences between murine and human dissections, mouse models can provide considerable insight and have gained in popularity owing to their reproducibility. Our results for three classes of geometrically different false lumens show that thrombus forms and extends to a greater extent in regions with lower bulk shear rates. Dense thrombi are less likely to form in high-shear zones and in the presence of strong vortices. The present data-driven study suggests that the proposed model is robust and can be employed to assess thrombus formation in human aortic dissections.

## Introduction

Dissection of the thoracic aorta is a life threatening event; it is responsible for significant morbidity and mortality in individuals ranging in age from children to young and old adults alike. Dissection can result from blunt trauma, but it often associates with aneurysmal dilatation and heritable connective tissue disorders, including Marfan syndrome, familial thoracic aortic aneurysm, Loeys-Dietz syndrome, Ehlers-Danlos syndrome, and so forth^1^. These dissections often propagate within the medial layer and connect with the true lumen to form a false lumen within the aortic wall, which in turn can remain patent, become partially thrombosed, or be completely filled with thrombus.

Thrombus formation is the normal physiologic response to prevent significant blood loss upon vascular injury, but its role in different disease conditions can be negative. Clinical findings increasingly suggest, for example, that patients with a partially thrombosed dissection are at a higher risk of rupture^2^. Conversely, completely thrombosed dissections have a better prognosis^3^. Indeed, it has been suggested that “complete thrombosis of the residual false lumen might be a sign of aortic wall healing and remodeling”^4^.

There has been a growing interest in the use of mouse models in vascular biology due to their availability and broad phenotypes, especially compared with other animal models. Most notably, chronic infusion of angiotensin II (AngII) in male apolipoprotein-E null (*ApoE*^-/-^) mice yields a reproducible model of dissecting aortic aneurysm, which often includes a false lumen with intramural thrombosis^5^. Although medical imaging now enables us to quantify the subject-specific geometry of the false lumen and to construct associated computational fluid dynamics models in both human and murine dissections^6–9^, no existing model can predict the development, growth, or arrest of an intramural thrombus. The present data-driven study is significant for it elucidates, for the first time, hemodynamic conditions under which an intramural thrombus forms in aortic dissections.

Quantifying continuum-level hemodynamics is fundamental for many reasons^10, 11^: circulating blood is the source of, amongst other components, the platelets, fibrinogen, and plasminogen that enable the thrombus to develop and subsequently remodel; increased fluid shear stresses can activate platelets; increased residence times within regions of low flow can promote platelet aggregation; and high shear rates can limit thrombus expansion into the flow field. Including the effects of platelets is particularly important in any numerical model that targets thrombus formation. Whole blood is commonly considered an incompressible Newtonian (or non-Newtonian) fluid in large arteries; this leads to continuum fields for blood velocity and pressure. The concentration of blood borne biomolecules can be resolved similarly using advection-diffusion-reaction equations, which provide spatio-temporal information on the concentration of, for example, fibrinogen (the precursor to fibrin) and plasminogen (the precursor to plasmin). Individual platelets or groups, in contrast, can be treated as Lagrangian particles. Building on our recent study^12^, we develop a particle-continuum model based on a force coupling method (FCM) that provides a flexible platform for *two-way coupling* of platelets (treated as semi-rigid spherical particles) and the background continuum flow. As a result, thrombus size and shape is affected by the local hemodynamics, particularly fluid stresses. Further, it is possible to introduce porosity to the forming thrombus by adjusting the radius of influence of each particle on the fluid as discussed in “Methods”.

While continuum-level hemodynamics plays a crucial role in thrombus formation and its final size and shape within a false lumen, the cellular and sub-cellular-scale hemostatic processes (*e.g*., platelet adhesion, aggregation and coagulation kinetics) cannot be ignored and must be taken into account in any multiscale model (see *e.g*.,^13–15^). We propose here a data-driven mulitscale numerical approach to address thrombus formation in dissecting geometries, whose length scales are significantly larger than microscopic scales encountered at the cellular levels. An additional challenge in modeling thrombosis is the long physiologic time-scales in the evolution of intramural thrombus, and its subsequent remodeling within the false lumen. Here, we take advantage of the apparent time-scale separation in this biological process that decomposes it into two main sub-processes, namely *acute thrombus formation* and the subsequent *growth and remodeling* of thrombus and the remnant wall.

## Results

Consider, first, the computed flow fields within each of the three lesions (Figure 1) with thrombus removed numerically. Results are represented as flow streamlines at peak systole, and without particles, to understand the pattern and strength of vortices that might contribute to thrombus deposition. The patterns of the vortical zones differ markedly by lesion and are affected mainly by the size of the opening into the false lumen. The small orifice in the small false lumen (Figure 1c) leads to the smallest vortex in the false lumen, whereas the orifice of the medium false lumen (Figure 1b) allows the circulation zone to grow in size. More notably, the large orifice size in the large dissection (Figure 1a) causes the flow to break into two significantly larger vortices inside the false lumen close to peak systole. Note, therefore, that Philips *et al*. analyzed λ_2_ distributions to detect regions of vortical flows inside the same lesions^16^. They showed that the negative λ_2_ regions form initially close to the proximal end of the false lumen opening and persist inside the false lumen for large parts of the cardiac cycle, consistent with results shown in Figure 1.

**Figure 1.**
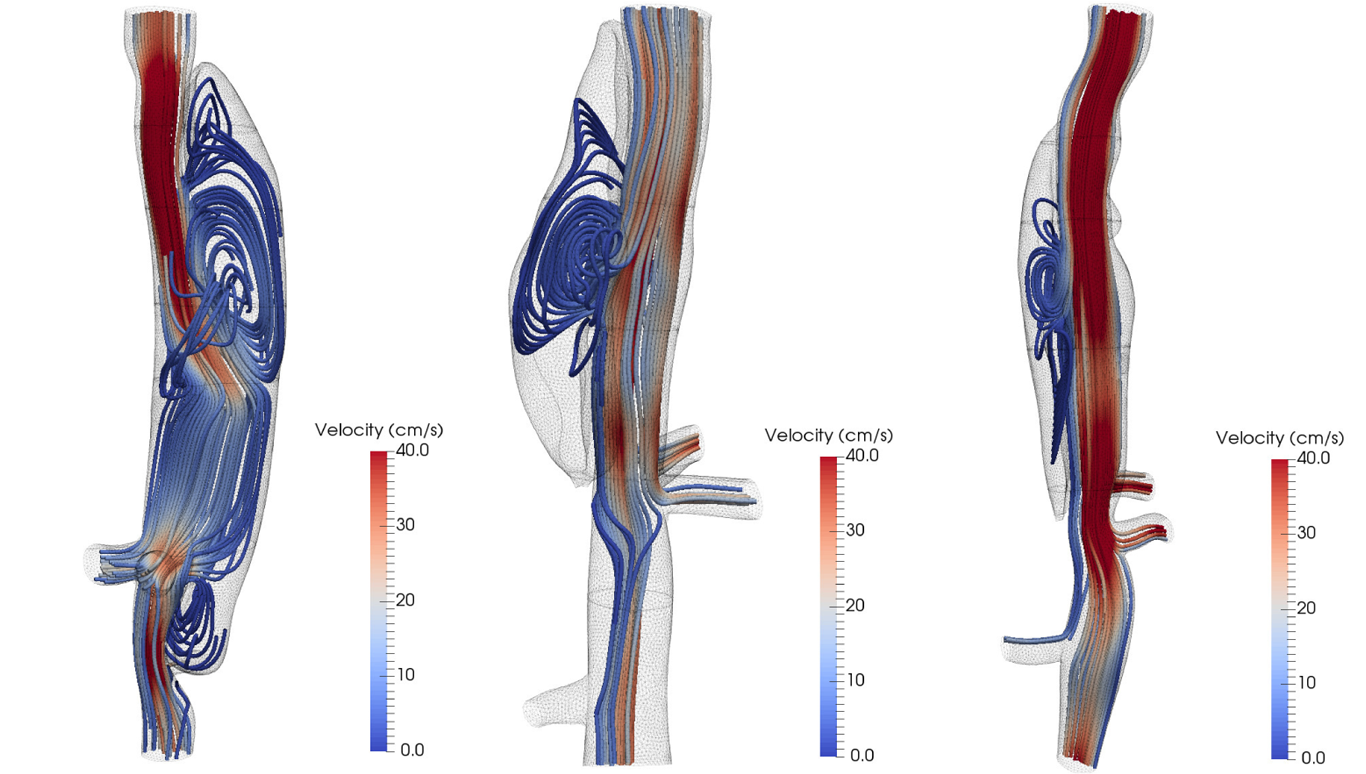
Snapshots of flow streamlines colored by vorticity magnitude at peak systole for the three lesions (blood flow is from top to bottom in all cases).

Figure 2 shows snapshots of particle distributions in the false lumen, where adhered particles in the thrombus are slightly larger than other particles. The streamlines also show a major vortex close to the orifice, which creates a hydrodynamically active region with significantly less particle adhesion and thrombus deposition. Continuum FCM volume fractions were calculated based on the adhered pseudo-particle positions using the FCM Gaussian kernel given in Eq. (10). As will be shown later (based on volume fractions φ_*fcm*_), our numerical simulations with FCM particles suggest that the size and shape of these vortices are significant determinants of final thrombus deposition inside the false lumen.

**Figure 2.**
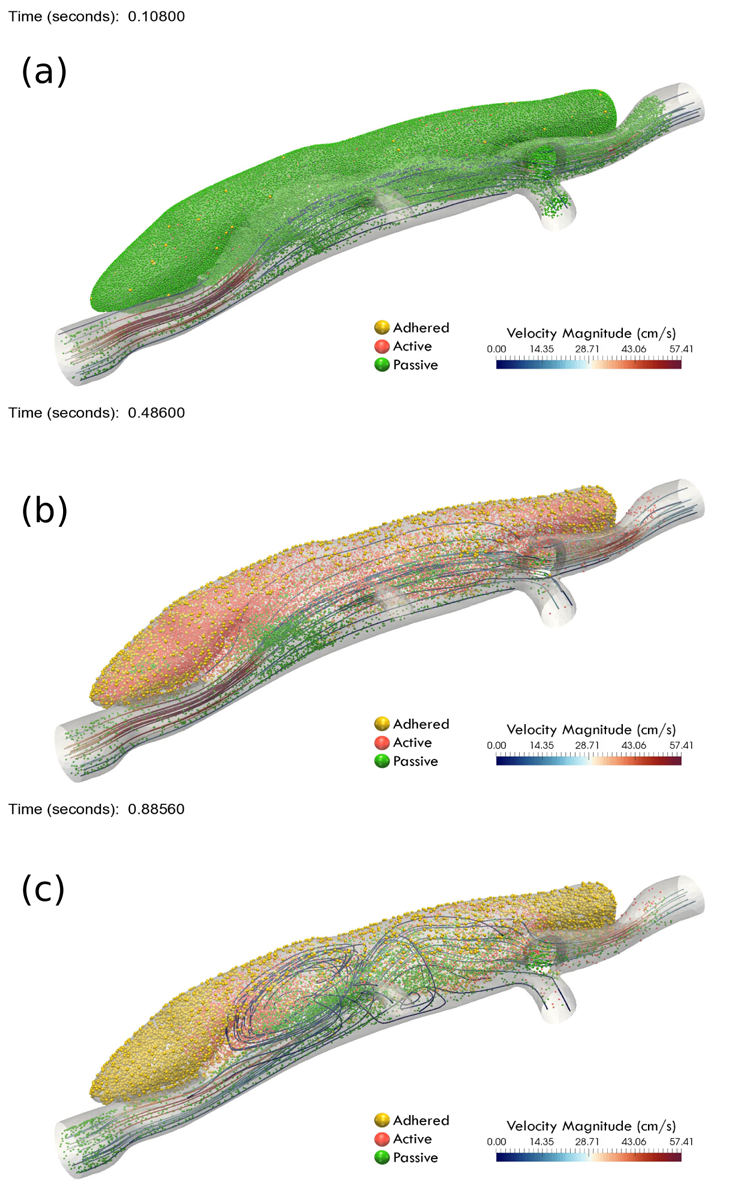
Snapshots of particle distributions in the large false lumen at three different time points of the simulation: (a) initial particle distribution; (b) intermediate stage at end diastole; (c) final stage at peak systole, along with a small number of representative flow streamlines colored by velocity magnitude. Particle color codes are green: passive particles; red: active or triggered particles; yellow: active and adhered particles. Not all the particles are shown for clarity, and flow is from left to right.

### Thrombus size and shape

#### Parameter estimation and sensitivity analysis

Two main parameters that affect thrombus deposition within the false lumen are the strength of the adhesive forces *D_e_* and the effective particle radius *r_eff_*. Whereas the value of *r_eff_* directly affects the cost of the simulations, since a lower effective radius implies a significantly higher number of particles 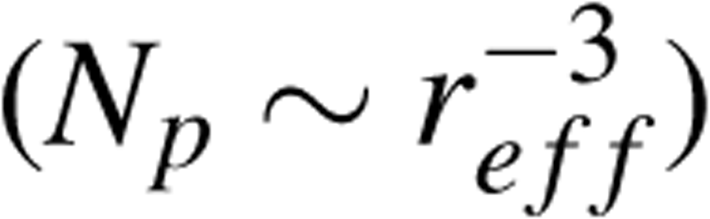, it also improves the predicted final shape that is otherwise driven by *D_e_*.

Noting that the clot volume fraction is a heterogenous spatial field (φ≡φ(*x,y,z*)), we quantified the variation of φ along the primary flow direction (*z*−axis) by integrating it on each planar cross section, namely 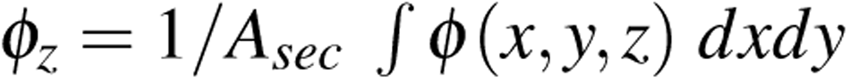, where *A_sec_* is the area covered by thrombus in each cross section. First, consider the evolution of clot volume fraction within the false lumen in Figure 3a for *D_e_* = 2 × 10^−3^ and *r_eff_* = 20*r_p_*. The results show an initial sudden increase in φ_*z*_ followed by a gradual increase towards its quasistatic distribution at *t\** = 100, which is equivalent to approximately 10 cardiac cycles, or *t* = 1 *sec*. Further, φ_*z*_ is plotted for three different values of *r_eff_* = 20*r_p_*, 30*r_p_*, 40*r_p_* in Figure 3b for comparison. The distributions of φ_*z*_ show a qualitative agreement for different effective radii at the two ends of the lesion (*i.e., z* > 1 and *z* < −1), whereas the results show significant deviations close to the orifice of the false lumen. An effective radius *r_eff_* = 40*r_p_* overestimates the shape of the thrombus at the opening, whereas the average volume fraction for *r_eff_* = 30*r_p_* and *r_eff_* = 20*r_p_* does not clearly show which one of the effective radii predicts better the thrombus size and shape; hence, a detailed analysis of thrombus shape similar to Figure 4 is required.

**Figure 3.**
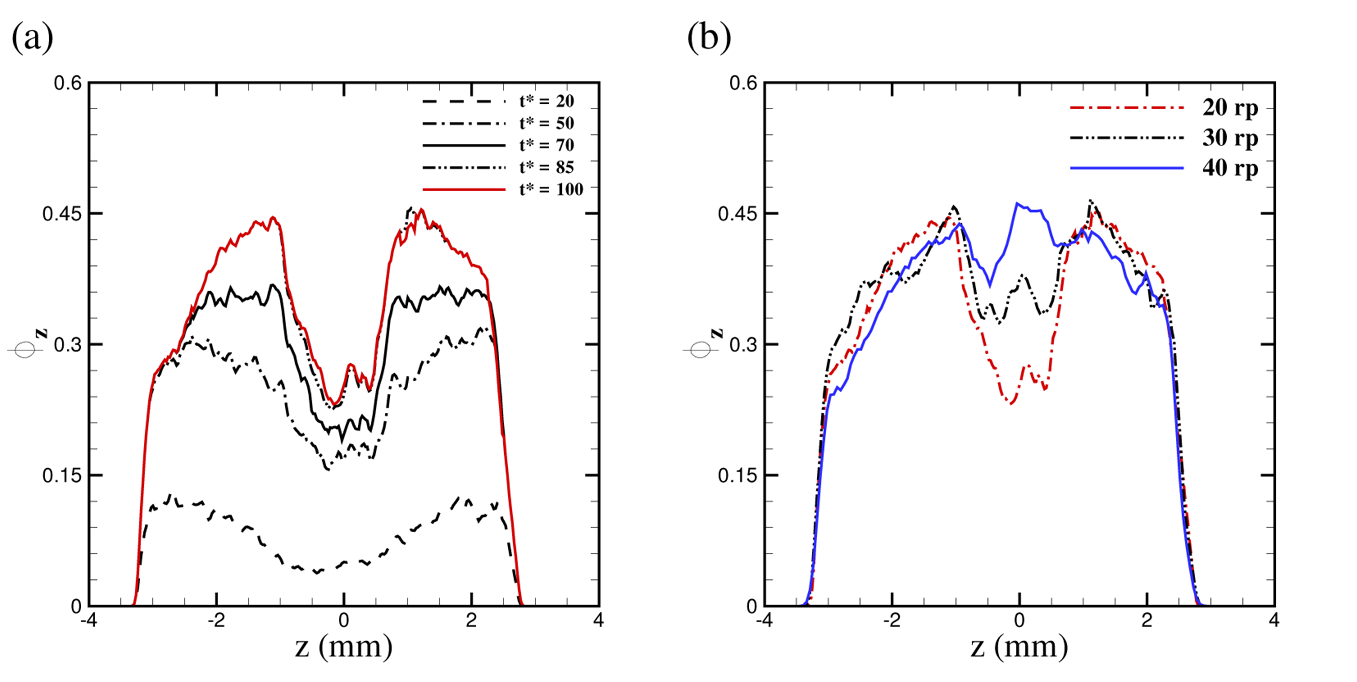
Average volume fraction of the clot φ_*z*_ computed in each cross section along the flow direction in the medium false lumen. (a) Temporal simulations suggest that clot volume fraction stabilizes after 10 cycles. (b) Simulations for different values of particle radius *r_eff_* suggest that smaller particle radius is more effective in capturing thrombus size and shape.

**Figure 4.**
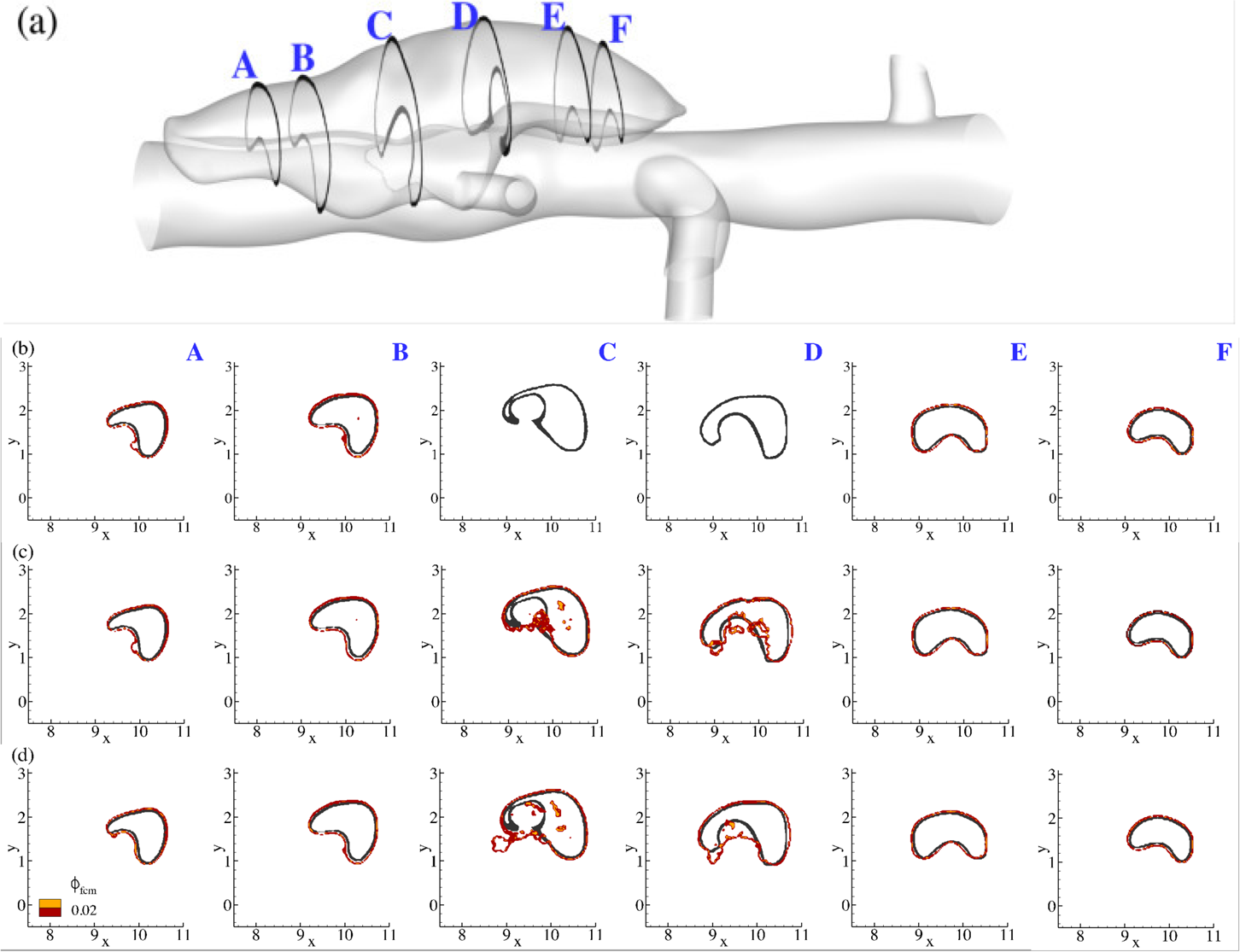
Effect of the adhesive force parameter *D_e_* on the size and shape of the clot (shown as an enclosed contour) for the medium false lumen for a fixed value of *r_eff_* = 30*r_p_*, where: (b) *D_e_* = 2 × 10^−4^; (c) *D_e_* = 2 × 10^−3^; (d) *D_e_* = 5 × 10^−3^. Comparison at 6 different cross-sections along the flow direction (left to right, or A-F), where dark contours show the segmented thrombus boundaries and red contours are the simulation results. Thrombus boundaries were estimated from simulations for threshold values of 0.02 < φ*_fcm_* < 0.04.

The effect of adhesive force coefficient *D_e_* is plotted in Figure 4a-4c for particles with radius *r_eff_* = 30*r_p_*, recalling that we seek agreement between the simulated clot (defined by the red boundaries in Figure 4) and the experimental thrombus from the OCT images (defined by dark boundaries in Figure 4). Here, we focus on the lesion with the medium false lumen and vary *D_e_* in the range of 2 × 10^−4^ ~ 5 × 10^−3^ (nondimensional units), while keeping *r_eff_* = 30*r_p_* (Figure 4a-4c). The best agreement was achieved for *D_e_* = 2 × 10^−3^ (except for the overestimation close to section C) as seen in Figure 4c-4d. Reducing *D_e_* by an order of magnitude (shown in Figure 4b relative to Figure 4c), caused the FCM to perform poorly in regions where the flow field is highly active with strong vortices (sections C and D in the figure), though all simulations captured thrombus size and shape very well at the two ends of the false lumen (sections A and F in the figure) since the flow dynamics is rather weak there. A further increase in *D_e_* to 5.0 × 10^−3^ overestimated the size and shape of the thrombus in the active vortical zone (sections C and D of Figure 4c) where adhesive forces are relatively high. Our simulations for all three lesions suggest that an adhesive force coefficient *D_e_* = 2 × 10^−3^ is appropriate for modeling the adhesion of pseudo-particles.

In addition to the choice of adhesive force, the effective radius of the activated particles plays an important role in determining the size and shape of the thrombus in the false lumen. As suggested by Figure 3, the value of *r_eff_* = 30*r_p_* overestimated the thrombus volume fraction and size especially in regions of higher fluid shear stresses. Hence, we used *r_eff_* = 20*r_p_* in our simulations for all three lesions, which leads to better prediction in thrombus shapes.

#### Acute evolution of thrombus size and shape

Next, we consider the evolution of thrombus deposition in the medium false lumen in Figure 5b, noting that thrombus formation is accelerated in our simulations within approximately 10 cycles due to increased size of an activated particle’s effective radius. By monitoring different cross sections along the false lumen, we observe faster thrombus growth and increased volume fractions closer to the ends of the false lumen, where φ_*fcm*_ can reach as high as 90%. Closer to the orifice of the false lumen (sections C and D), however, simulations predict a partial thrombosis with a lower volume fraction, indicating a less dense thrombus close to the remnant wall.

**Figure 5.**
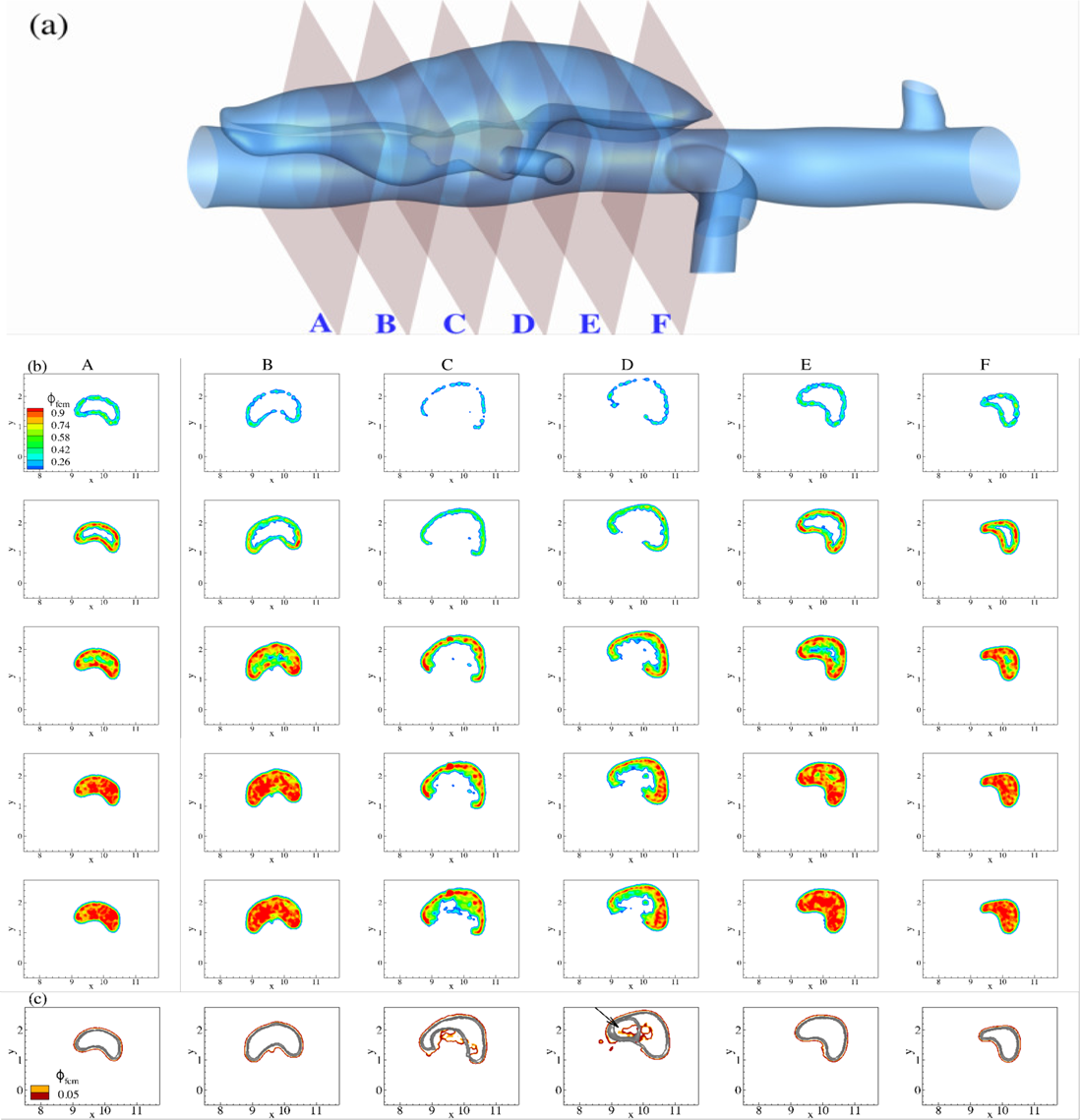
(a) Guide for planar cross-sections in the medium false lumen taken along the flow direction (along +*z* axis with flow from A to F). (b) Time evolution of continuum **FCM** volume fraction φ*_fcm_* field for 6 cross sections labeled as above. First row is plotted after the first 2 cardiac cycles, with time incremented 2 cardiac cycles for every subsequent row. (c) Cross-comparison of simulated thrombus boundaries (red) with experimental shapes (gray) extracted from OCT images.

We performed an analysis similar to Figure 3 to identify when a quasi-stable thrombus shape had been reached in our simulations. Once the size and shape stabilize, we extracted the boundaries of the thrombus and compared them with thrombus geometries segmented from OCT images in^16^ (see Figure 5c for the medium false lumen). We achieved very good agreements in less hydrodynamically active zones (sections A, B, E and F). In more active regions, the agreement was acceptable given the uncertainty in both the simulation parameters and experimental measurements (*e.g*., OCT images have spatial resolution ~ 7µ*m*). Most notably, the boundary of the false lumen in section D, which forms a ring (shown by an arrow in Figure 5c), was captured in our simulation despite a slight offset in the location of the simulated false lumen relative to the experiment.

Our simulations suggested a strong correlation between hemodynamic forces and final thrombus size and shape. To further examine this finding, we performed FCM simulations for two additional simulated lesions with different orifice sizes that created different hemodynamic environments (cf. Figure 1). We present the evolution of thrombus formation for the small false lumen in Figure 6b, for which the vortical structure close to the orifice is smaller than that for the medium false lumen (although vorticity magnitudes are higher) as shown in Figure 1c.

**Figure 6.**
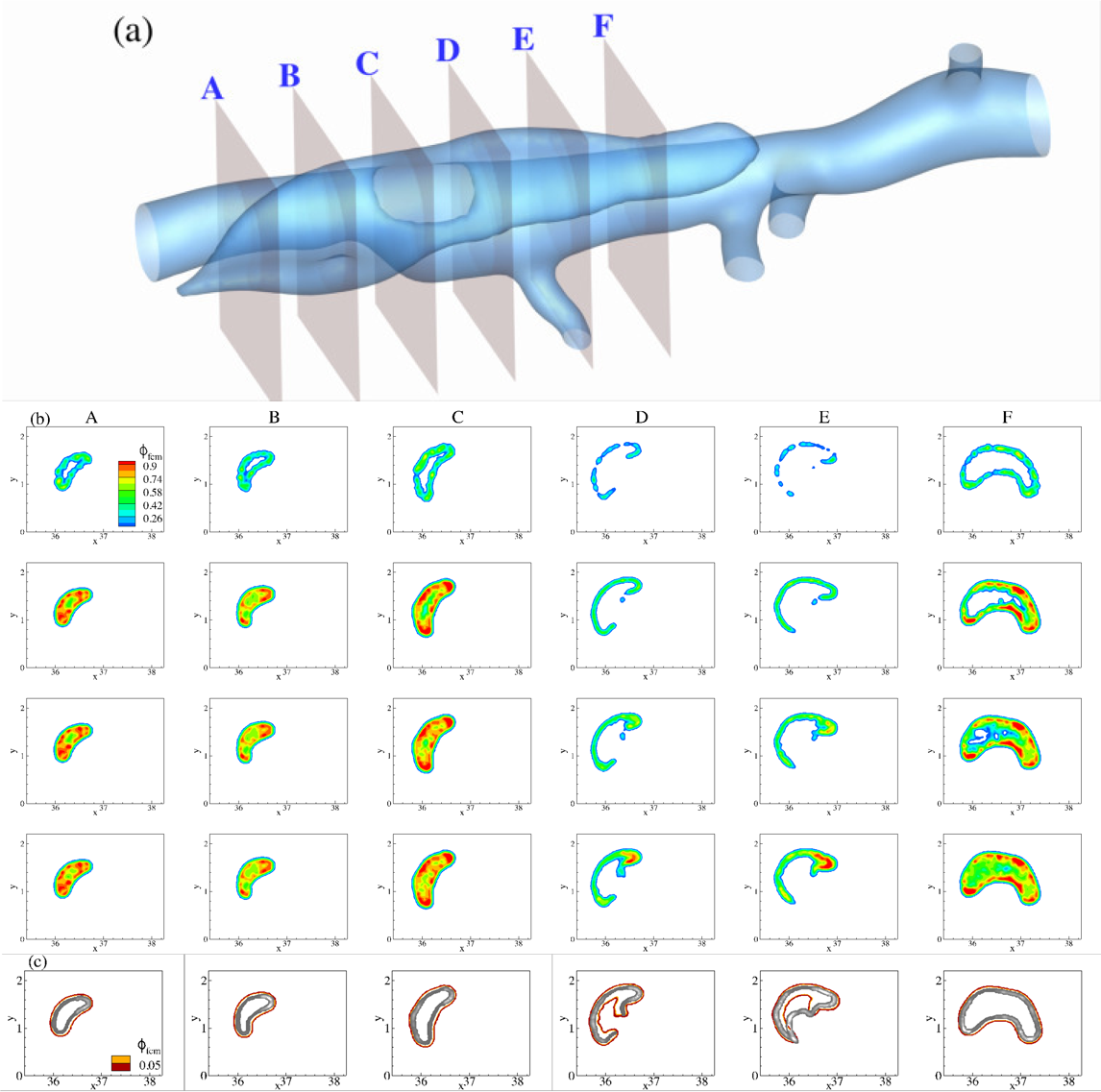
(a) Guide for planar cross-sections in the small false lumen taken along the flow direction (along −*z* axis with flow from A to F). (b) Time evolution of continuum FCM volume fraction φ*_fcm_* field for 6 cross sections labeled as above. First row is plotted after the first 2 cardiac cycles, with time incremented 2 cardiac cycles for every subsequent row. (c) Cross-comparison of simulated thrombus boundaries (red) with experimental shapes (gray) extracted from OCT images.

Note a fast growth of thrombus closer to the ends of the false lumen (*i.e*., sections A-C and E-G), with the size of the patent lumen reduced significantly and limited to the vicinity of the orifice (section D). Taking into account the pattern of vortices in that region (Figure 1c) and the direction of flow (along −*z* axis), the effects of hemodynamics on the final size and shape is prominent. Further comparison of the final size and shape between simulation and experiment shows a good agreement in most regions of the false lumen, albeit with a slight underestimation in the size of thrombus at cross-section E. Close to the orifice, however, thrombus formed adjacent to the false lumen wall (section D), as predicted by the simulation.

Next we performed FCM simulations for the largest false lumen, which also had the largest orifice among the three lesions. The wider opening into the false lumen creates a significantly larger vortex than the other two lesions, which breaks into two strong vortices close to peak systole (cf. Figure 3b and supplementary video). These vortices, and the resulting shear forces, prevent platelets from aggregation and forming a firm and stable thrombus inside the false lumen along most of the width of its opening. This can be clearly observed from the evolving thrombus shape in Figure 7b along sections B to E. The two ends of the false lumen form more stable thrombi, however, similar to other lesions due to weaker hydrodynamic forces present in those regions.

Comparison with experimental thrombus shape in Figure 7c shows an under-prediction of thrombus size at sections B and D. Section D lies at the center of the false lumen, where the two counter rotating vortices meet during peak systole and, thus, form a stagnant region. This could potentially result in a higher accumulation of platelets and their residence time in that small region depending on the initial distribution of platelets and other coagulants in a dissecting aneurysm, which is not exactly known *a priori*. Further, we believe that the smaller size of thrombus close to section B (shown by the arrow in Figure 7c) is due to significantly higher shear forces in that region compared to other lesions.

**Figure 7.**
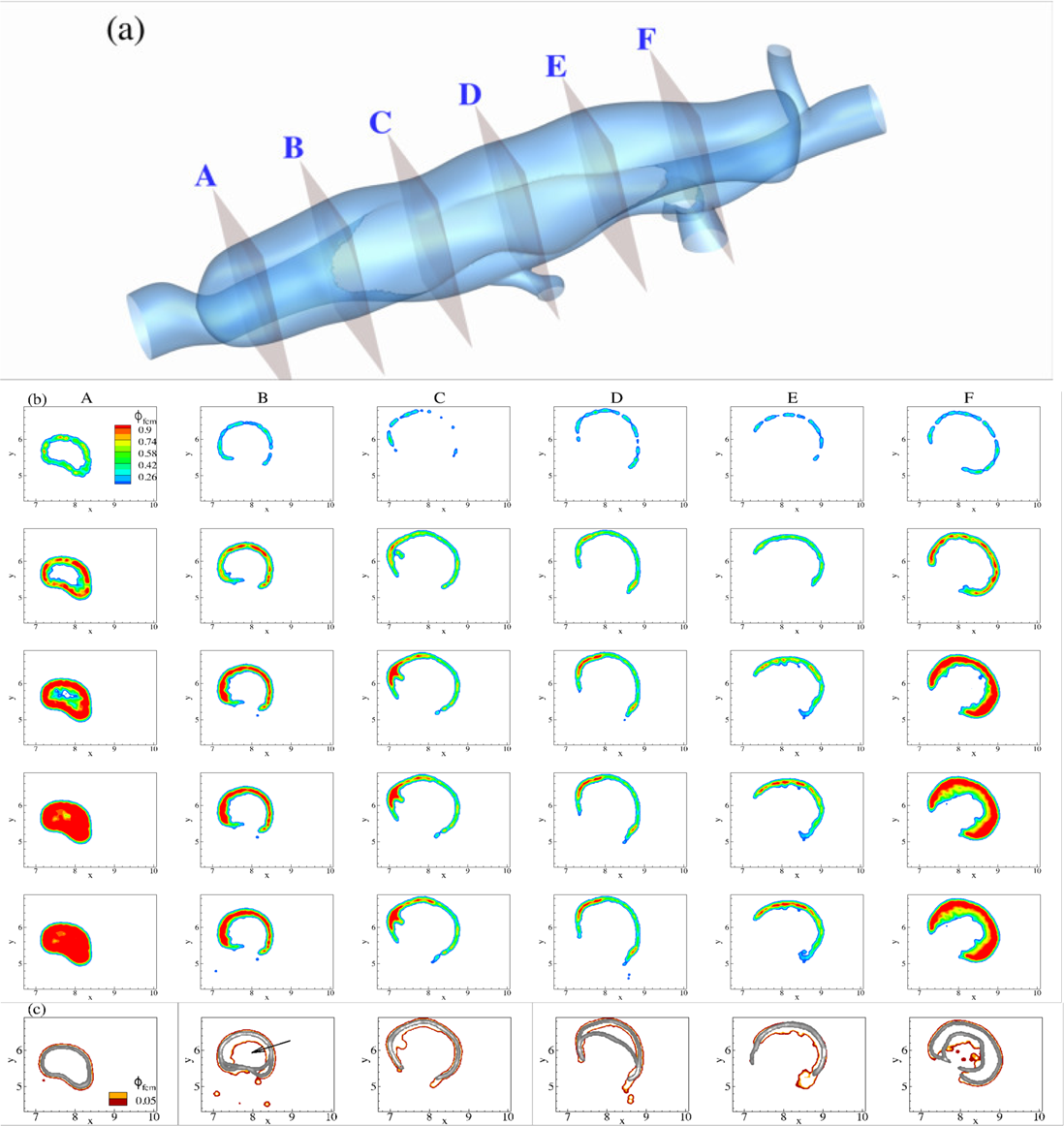
(a) Guide for planar cross-sections in the large false lumen taken along the flow direction (along −*z* axis with flow from A to F). (b) Time evolution of continuum FCM volume fraction φ*_fcm_* field for 6 cross sections labeled as above. First row is plotted after the first 2 cardiac cycles, with time incremented 2 cardiac cycles for every subsequent row. (c) Cross-comparison of simulated thrombus boundaries (red) with experimental shapes (gray) extracted from OCT images.

To further clarify the role of shear rates on the progression of thrombus deposition, we simulated blood flow within the false lumen at two time-points: the onset of dissection and after thrombus deposits. The results for scalar shear rate 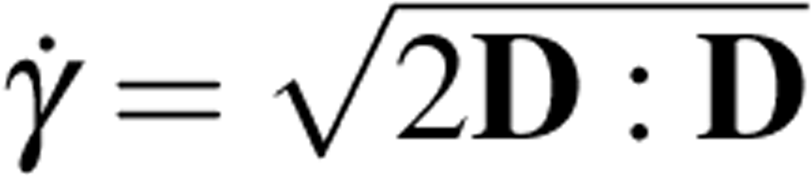 are shown in Figure 8 for large and medium false lumens, where **D** = (**∇u** + **∇u^T^**)/2 is the symmetric part of the spatial velocity gradient.

**Figure 8.**
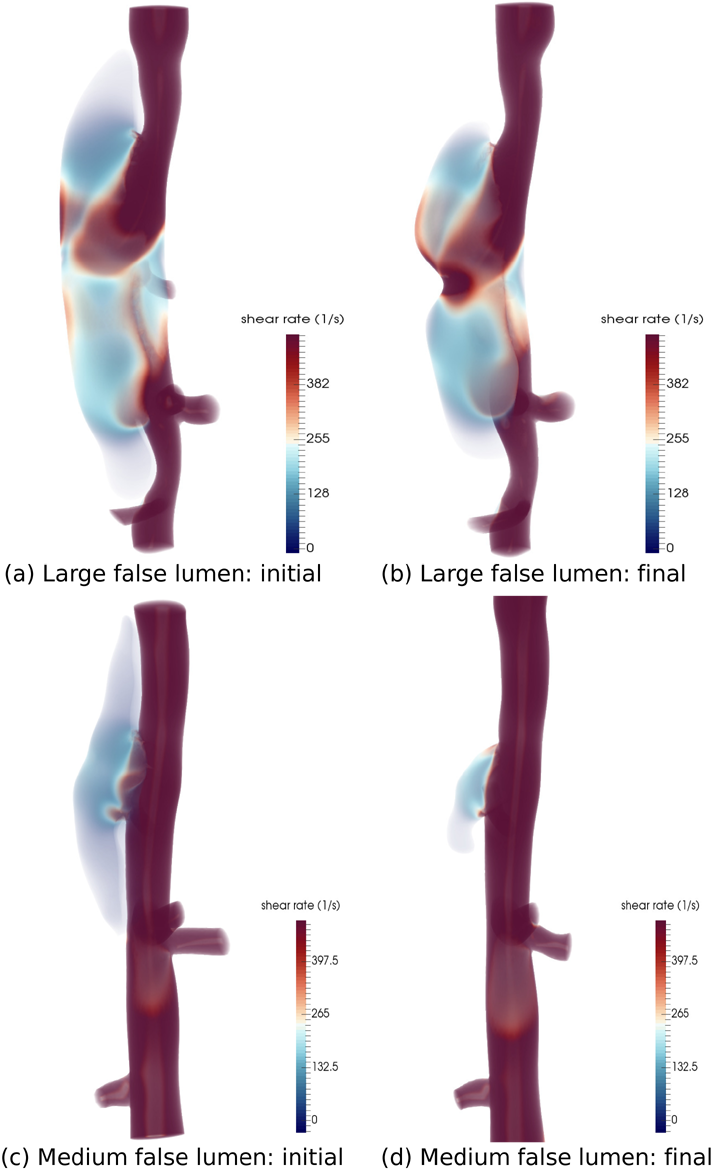
Volume rendering of scalar shear rate 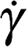 at peak systole for two lesions at the initial (left) and final (right) simulation times (blood flow direction is from top to bottom in all cases): (a), (b) for large false lumen; (c), (d) for medium false lumen.

The results show clear distinctions in the magnitude and location of high shear rates within the false lumen between large and medium lesions: higher shear rates (≈500 *s*^−1^ are present at peak systole inside the large false lumen, covering a wider region inside the false lumen, whereas shear rates are close to 100 *s*^−1^ in the medium false lumen (see Figure 8a and 8c). One common pattern between these lesions is relatively weak shear rates near the ends of the false lumen, which are presumably sites where thrombus starts depositing. As thrombus progresses toward the center of the false lumen, it is subject to increasingly higher shear rates, which eventually prevent further thrombus formation. Hemodynamics simulations for the experimentally observed false lumens show distinct flow regimes and shear rates in these two lesions (see Figure 8b and 8d). The shear rates remain relatively low, close to 100 *s*^−1^ inside the final shape of the medium false lumen. In contrast, shear rates increase further inside the large false lumen, especially in the vicinity of the thrombus surface, thus preventing further thrombus formation and suggesting why the large false lumen remains nearly patent.

### Transport of fibrinogen and platelets in the false lumen

Knowledge of local concentrations of fibrinogen (Fbg) and platelets inside the false lumen is essential to estimating the composition of thrombus. We performed pilot studies for the transport of Fbg and platelets inside the false lumen by solving the advection/diffusion Eq. (9). We set the normalized concentration of both species equal to 1 everywhere and performed simulations with zero flux boundary conditions for long periods (~ 10 cardiac cycles) to reach stable distributions inside the false lumen. Results for the medium and large false lumens are shown in Figure 9.

**Figure 9.**
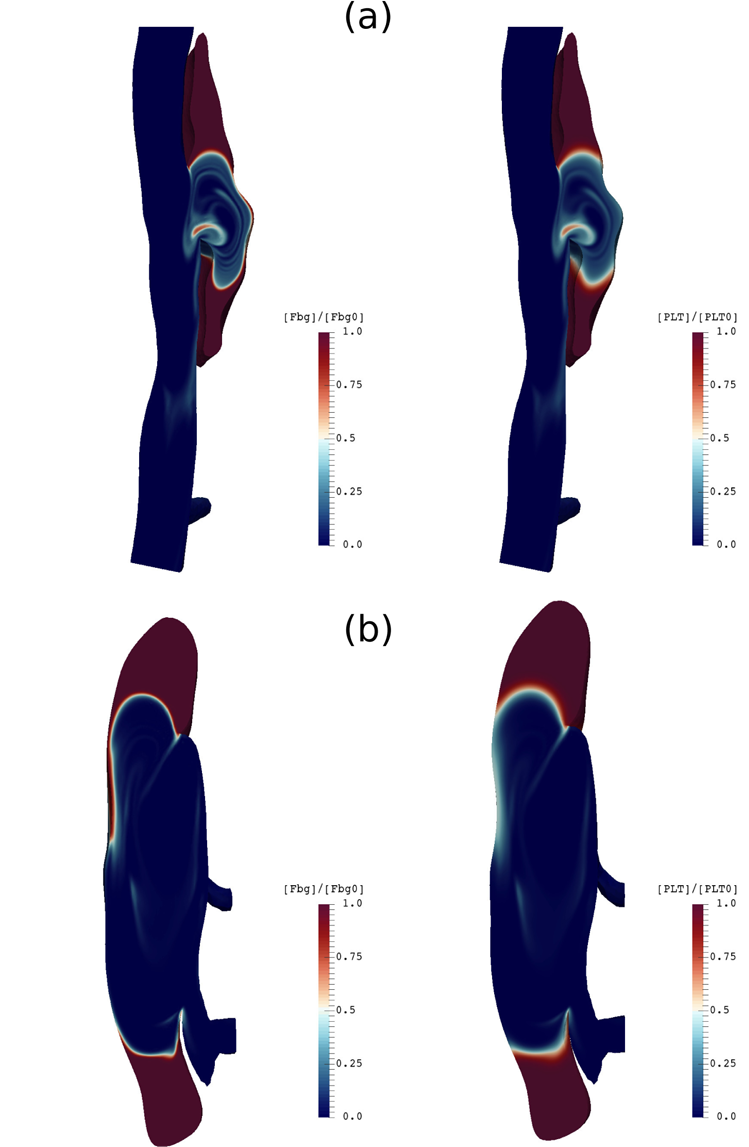
Contours of normalized Fbg concentration for (left) and platelet concentration (right) plotted on a longitudinal slice along the flow direction (top to bottom) for (a) medium and (b) large false lumens after 10 cardiac cycles.

Distributions show a similar pattern for Fbg and platelets for both medium and large false lumens in which the concentrations remain less affected by the flow close to the ends of the false lumen, where bulk shear rates are lower. At the center of the false lumen close to its opening, however, where shear rates are higher, we observe large regions depleted of Fbg and platelets. Interestingly, the depleted regions co-localized with regions of high bulk shear rates (see Figure 8) and, thus, could be used as a guide for the extent of acute thrombus growth and the size of the final false lumen.

## Discussion

Our subject-specific numerical results, based on three geometrically different lesions, suggest that the initial size and the shape of the false lumen and the size of its orifice are highly crucial in dictating the progression of thrombus formation inside the dissection. The salient feature of all three lesions is enhanced accumulation of thrombus at the ends of the false lumen, which associate with significantly lower local shear rates (≪ 50 *s*^−1^). Closer to the opening of the false lumen, however, stronger vortices and higher shear rates prevent further thrombus growth. These results suggest a threshold shear rate value below which thrombus is more likely to form, 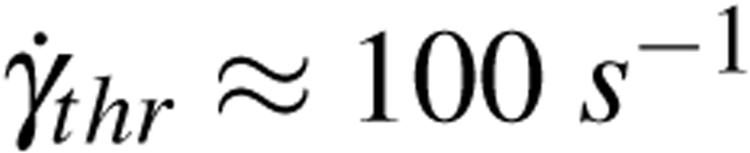. A more rigorous statistical analysis with more subjects would be needed to confirm this apparent threshold value, however.

It is acknowledged that murine aortic dissections differ from human dissections in terms of location, geometry, and extent of the false lumen. Human dissections often manifest as longitudinal false lumens that are typically very long, and may develop multiple proximal and distal tears. The Reynolds number of the hemodynamics are higher in the false lumen, which can cause secondary flows and different flow patterns. Numerical simulations of thrombus formation in patient-specific aortic dissections are relatively rare. In a recent study, Menichini *et al*.^17^ proposed a thrombosis model for predicting the false lumen in patient-specific type B aortic dissections. The model is based on continuum fields of activated and resting platelets as well as most relevant coagulation enzymes. Although proposed for human dissections, the results are similar qualitatively to the present murine results in a sense that thrombus formation initiates in regions of low wall shear stress or equivalently low bulk shear rates.

The multiscale numerical method we proposed here is novel in that it tackles both short-term and long-term processes (see Figure 10), noting that the present study addressed only the first portion of the numerical approach, namely the modeling of initial thrombus deposition until a quasi-static shape has been reached. Clearly, the challenge in this problem is long-term simulation of particles actively coupled with flow. The complexities in geometry and flow conditions and the large size of aortic dissections may require hundreds of thousands of FCM particles to represent platelets, which imposes a restrictively high computational cost for such simulations in large domains. To overcome this difficulty, we initially distribute particles in the false lumen with thrombogenic surfaces and let the flow transport the particles and their aggregates. Further, the increase in the size of particles upon activation is crucial in reducing the number of platelets in the domain and, hence, accelerating the process of thrombus formation. These numerical treatments altogether, enable us to address the short-term process as a *final-value* problem instead of simulating the original initial-value problem.

**Figure 10.**
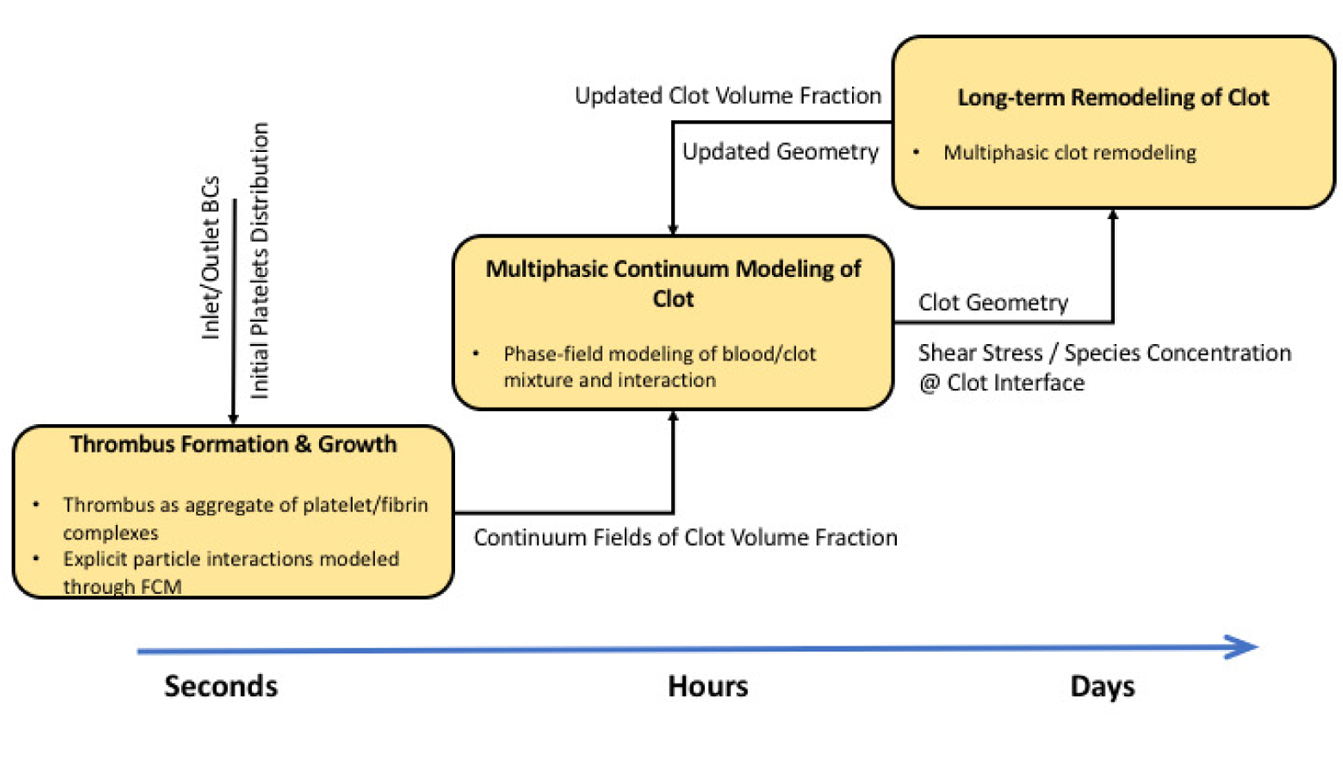
The proposed multiscale numerical model of thrombus formation and growth/remodeling of the remnant wall in a
dissecting aneurysm. The platelet activation/aggregation model, based on FCM, is the most expensive module in this framework and, thus, is only used to model the formation and propagation of the clot for several seconds after dissection. Once a stable intramural thrombus is formed, we transform the platelets Lagrangian distribution in the aggregate clusters into continuum fields of clot volume fraction, which can be subsequently used in a phase-field numerical approach to model interaction of blood with thrombus, and its further remodeling.

Long-term remodeling of the thrombus based on the constrained mixture theory^18^ will require a continuum representation of the thrombus shape and its composition. Therefore, it is desirable to represent the thrombus as a separate continuous phase field. Volume (or mass) fraction of clot constituents can be estimated using the FCM volume fraction (φ_*fcm*_) and local platelet and fibrinogen concentrations. Rheological studies have shown that fibrin clots exhibit viscoelastic behavior^19^, which will allow us to model the clot-blood flow interaction using a phase-field approach and a multiphase formulation^19^. Further, the transport of other relevant biochemical concentrations to fibrinolysis (*e.g*., plasminogen) may be modeled in blood (single-phase) and within the thrombus (multiphase) through a multiphasic approach. We will address the issue of long-term remodeling of thrombus and its interaction with blood in a separate study.

In the present study, we were mainly interested in the acute size and shapes of the false lumen, thrombus, and thus initial rather than the long-term evolution. We accelerated the process several-fold by (i) distributing particles uniformly inside the initial false lumen assuming that blood enters the false lumen upon onset of dissection and platelets are almost everywhere, (ii) assuming that all dissected mural surfaces expose the blood to damaged collagen and are thus thrombogenic. Hence, platelets in the vicinity of the wall become activated and can adhere to the walls; once activated, they grow up to 20 times larger by turning into pseudo-particles that bear the physiologic density of platelets and concentration of fibrin. Clearly, the presence of activated platelets is essential in the progression of thrombus formation. Shear-induced activation of platelets due to an accumulated stress history can also stimulate platelets in high-shear flow conditions, such as blood flow through atherosclerotic plaques and mechanical heart valves^20, 21^, or even abdominal aortic aneurysms^22–24^. We did not consider shear history since we felt that biochemical effects dominate with wall damage. Nevertheless, Menichini *et al*.employed a mechanical activation model to incorporate the shear stress history of platelets along their trajectories and found that the highest activation levels occur for platelets crossing the entry tear when transported into the false lumen. The time-averaged wall shear stress close to the entry tear in their human study was about 20 *Pa*, which is significantly higher than values in the present study of murine dissection (<1 *Pa*). Therefore, in our model, only resting platelets that are within a short distance to the damaged wall will be initially activated while other platelets will become activated via contact.

In the FCM method, inter-particle forces are transferred to the carrier fluid through smooth Gaussian kernels. Using the same kernels, we generated continuous fields of volume fractions for the distribution of particles. Volume fraction φ_*fcm*_ served two purposes in our study. First, it works as an indicator function as to where thrombus is more likely to form in the false lumen, which in turn, shows boundaries of the thrombi (*e.g*., Figure 4). Second, it identifies regions in which more platelets adhere and, hence, thrombus is denser (indicated by higher φ_*fcm*_ values) versus regions where shear rates are higher and less deposition occurs (lower φ_*fcm*_ values). The agreement achieved between the simulated and observed boundaries of the thrombus, despite uncertainties in numerical parameters and experimental measurements, confirms the general ability of our approach. Detailed analyses of the final shapes of false lumen show only slight deviations in a few particular regions such as slice D in the medium false lumen (Figure 5) and slices B and D in the large false lumen (Figure 7). The common feature for all of these regions is the locally high shear rates. Currently, our FCM model is calibrated for flow regimes inside the false lumen and presumably controlled by low-shear conditions. Therefore, particle adhesive forces are not adjusted in the regions of elevated shear rates, which may have lead to the mentioned deviations. A shear-dependent calibrated model (cf.^12^) could improve the numerical predictions further.

Because the FCM volume fraction cannot determine the composition of a clot in terms of its dominant constituents, we considered transport of the most important species that contribute to a clot, namely fibrinogen and platelets. Knowing local concentrations of these components along with φ_*fcm*_, and using an empirical correlation, we were able to estimate the volume fraction of clot through Eq. (11). For example, in experimental work by^25^, the fiber volume fraction was estimated by the empirical relation φ_*f*_ = *c_Fbg_*/[ρ_*Fbg*_ 0.015 log(*c_Fbg_*) + 0.13], where *c_Fbg_* (in *mg/mL*), is a key parameter, with ρ_*Fbg*_ = 1.4 *g/mL* the density of a single fibrinogen protein. Further, the volume fraction contribution of platelets φ_*p*_ can be directly evaluated based on each platelet’s volume (assuming a spherical particle of radius 1.5 micron) and the local platelet number density.

Solving the transport Eq. (9) is normally much cheaper than using particles coupled with the flow solver. Therefore, it is a straightforward task to perform parallel pilot simulations in the patent false lumen to estimate platelet and fibrinogen/fibrin concentrations that can be used in the evaluation of clot volume fraction. Further, it is noteworthy that the transport Eq. (9) for platelets can be used to estimate platelet residence times in a similar fashion. Evidence suggests that regions with higher residence times are more prone to initiate thrombus and further growth^17, 22^. Having accurate spatio-temporal measurements of clot volume fraction φ_*c*_ is essential for long-term growth and remodeling simulations. Multiphasic modeling of blood and clot interaction along with clot remodeling from an initially fibrin-dominated structure to a collagenous mass (cf.^26^) using a constrained mixture model will be important to simulate as well.

In conclusion, we present for the first time a particle-continuum numerical model to predict acute thrombus size and shape within aortic dissections. The numerical method is based on a two-way coupling of Lagrangian particle transport with blood, where a phenomenological model for inter-particle forces is used to mimic platelet adhesion and aggregation. Our numerical results show strong effects of hemodynamics on thrombus growth, and, thus, final shapes of false lumen. These results were previously reported in the experimental findings for three classes of murine suprarenal aortic dissections^16^. The availability of three different lesions was extremely important in the calibration of our model and in performing posterior analyses of numerical results. Numerical predictions of the location and extent of thrombus is satisfactory, and we are looking to extend the model to human dissections, which could be beneficial to clinical decision-makings in possible endovascular treatments.

## Methods

Aortic dissection was induced via a standard subcutaneous infusion of AngII at a rate of 1000 *ng/kg/min* in adult male *ApoE*^-/-^ mice. Multi-modal imaging data, obtained by combining 3D ultrasound and optical coherence tomography (OCT), were previously used to construct computational domains while pulsed wave Doppler and anatomic 3D ultrasound were used to estimate the average flow velocities at the inlet, outlet and four branches of the aorta^16^. Herein, we consider three classes of aortic dissection that did not rupture (Figure 11): one with a large false lumen with little thrombus, one with considerable thrombus but a modest residual false lumen, and one that is filled with thrombus. All experimental procedures were approved by the Institutional Animal Care and Use Committee (IACUC) at Purdue University and were carried out in accordance with the approved guidelines^16^.

**Figure 11.**
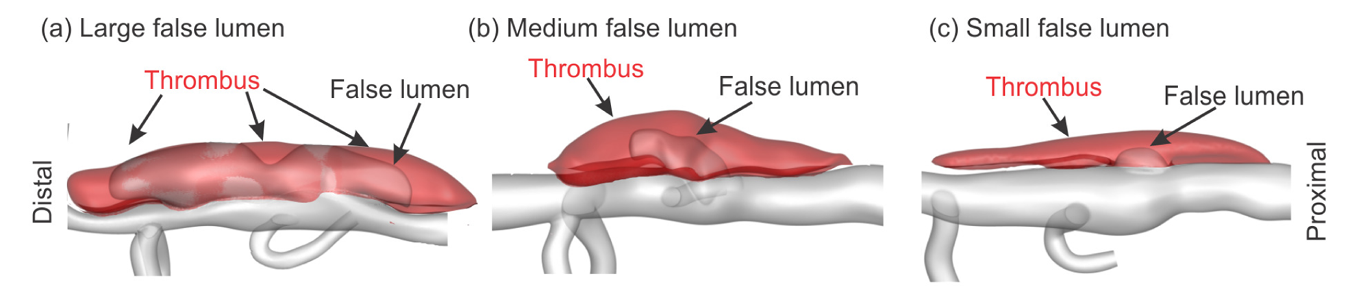
Reconstructed geometries from 3D ultrasound and OCT image stacks of three dissections of the suprarenal aorta in the *ApoE^-/-^* mouse; from^16^. The shaded red volume represents the thrombus whereas the poink volume is the false lumen with flowing blood; the grey shows the patent vessel. The direction of flow is from right to left, that is, from the proximal to the distal end.

### Particle-Continuum model

Due to the long-time evolution of intramural thrombus, formation, maturation, and remodeling within the false lumen, a multiscale approach is necessary. More specifically, it is best to decompose the entire process into two main parts: first, the initiation and formation of thrombus within the false lumen, which occurs on the order of minutes to hours following the dissection and, second, the growth and remodeling of the clot, which is mainly related to fibrin degradation and collagen production, which occurs on the order of days to weeks. In this paper, we focus on the former – the acute phase that establishes the initial size and shape of the clot. Toward this end, we focus on the platelet activation, aggregation, and formation of the initial fibrin mesh.

#### Modeling platelet transport and aggregation using FCM

The motion of platelets within a flow field and their adhesion to a damaged intramural dissection surface are solved together by coupling a spectral/hp element method (SEM)^27^ with a FCM similar to the work of Pivkin *et al*.^28^. Specifically, SEM is used to solve the flow field on a fixed Eulerian grid, whereas FCM is implemented to describe the two-way interactions between the blood flow and platelets. Due to the high concentration of fibrinogen in blood, and thus initially within a dissection, and fairly weak flow conditions inside the initial false lumen, the clot is considered to consist mainly of fibrin^29^. Fibrin monomers polymerize and form networks that capture platelets and other blood cells to form a loosely packed, porous clot. In our model, platelets are assumed to exist in three different states, namely *passive, triggered*, or *activated*, a model first proposed in^28^. In passive or triggered states, platelets have their physiological radius of *r_p_* = 1.5 µ*m* and are non-adhesive (Figure 12a). Initially, passive platelets are distributed uniformly inside the false lumen (Figure 12a). If a passive platelet interacts with an activated platelet or the damaged wall bounding the false lumen, however, it becomes triggered and switches to an activated state after delay time τ_*act*_ = 0.1 − 0.3 *s*. Upon activation, pseudo-platelets (*i.e*., platelets and associated fibrin) grow in size to an effective radius *r_eff_* (Figure 12b). This approach allows us to use fewer platelets than the physiologic concentration and to grow the size and shape of the clot once the platelets come in contact with the injured wall or each other.

**Figure 12.**
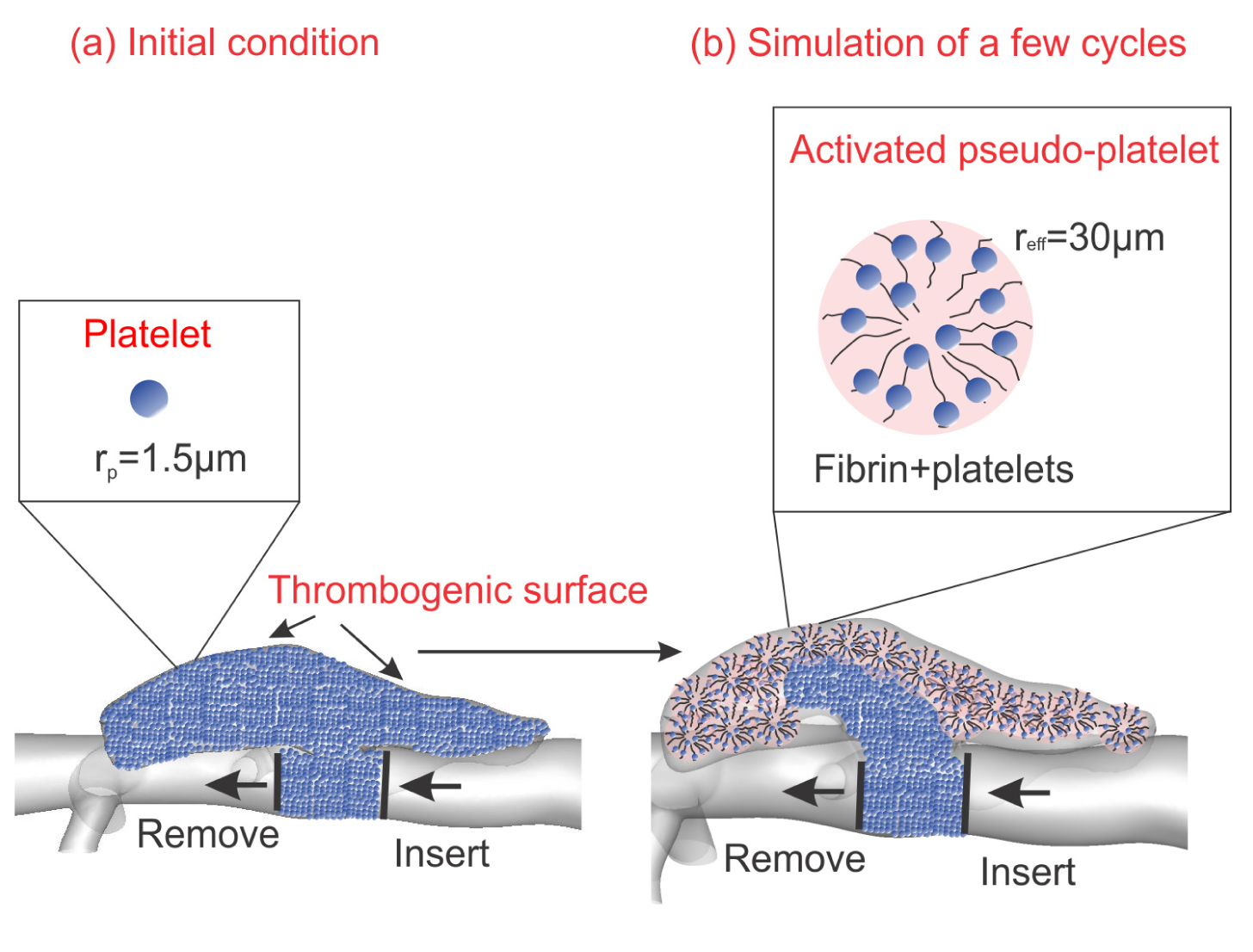
(a) Initial platelet distribution in the instantaneous false lumen, with the actual thrombus removed numerically. Also shown are locations along the artery where new platelets are inserted into the domain and outgoing platelets are deleted from the simulation. *Inset*: passive platelets as spherical particles of radius *r_p_* = 1.5 μ*m*. (b) Platelets change their size once they come into contact with the dissected wall and become activated, catalyzing the conversion of blood borne fibrinogen into semi-solid fibrin – *Inset*: activated platelets with increased effective radius *r_eff_* = 20.40*r_p_*.

The FCM has been described in detail previously^12^. Briefly, the translational velocity of each platelet (passive or activated) is estimated by the local average of the fluid velocity weighted by a Gaussian kernel function. The governing equations for the incompressible flow are the mass balance (continuity) and linear momentum balance (Navier-Stokes) equations

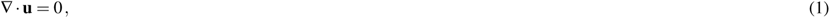

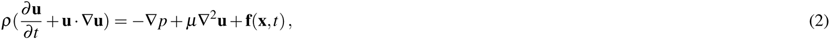

where u, *p*, and µ are the flow velocity, pressure and blood viscosity, respectively, with body force fused to capture the effects of the particles (platelets) on the flow. Specifically,

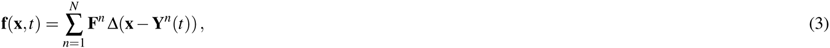

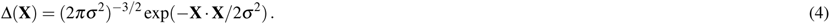

Here, F^*n*^ is the force due to particle *n* in the local region of interest. Moreover, the contribution of each platelet, whose center of mass is located at **Y**^*n*^, to the flow at position x is smoothed by a Gaussian distribution kernel Δ(**X ≡ (x − Y^n^**)) with σ the standard deviation of the kernel, which is related to particle radius *a* = (*r_p_* or *r_eff_*)through 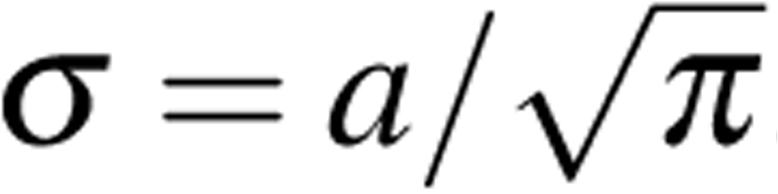. The governing equations are written in weak form and the domain is discretized using spectral elements that allow high order Jacobi polynomials. Time integration is performed using a semi-implicit splitting scheme^27^. The velocity of each platelet **V**^*n*^ is calculated by interpolating the local flow velocity at the location of a platelet using the same Gaussian kernel of Eq. (4) although different standard deviations may be used for force and velocity interpolations. Specifically,

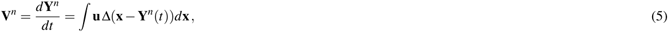

where position vectors for all the platelets are updated at each time step using a second-order Euler forward scheme. Further, the net force **F**^*n*^ acting on each platelet is written

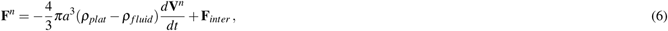

where the first term is the inertial force resulting from the difference in mass density between the platelets and blood flow and the second term accounts for interaction forces between platelets and the wall, which represent effects of different ligand-receptor interactions. Specifically, **F**_*inter*_ = −∂*U*/∂*r*, where *r* is the distance between platelet centroids and the potential *U* = *U_Morse_* + *U_exp_*.

Specifically, we use a phenomenological model based on Morse potential *U_Morse_*, having units of (Nm), to model attractive interactions between platelets, written as

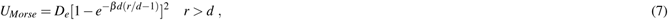

where *D_e_* is the energy depth contributing to the strength of the interaction force, β controls the width of the energy well, and *d* = 2*r_p_* (or 2*r_eff_* if activated) is the equilibrium distance between two platelets. To avoid instability due to strong Morse repulsive forces, we use an exponential repulsion potential in the form of

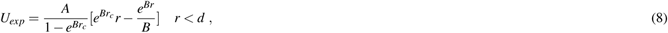

where the parameter *A* defines the maximum repulsive strength as *r* → 0, *B* is an adjustable parameter that defines the shape of the repulsive force (see Figure 13b), and *r_c_* is the cutoff radius. By setting *B* to a large positive number (here *B* = 25 in nondimensional units) and letting *rc* =*d*, we can obtain a hard-sphere repulsive potential. We set the parameter *A* such that the ratio of maximum repulsive to attractive forces is 5 (Figure 13b). Note that unlike the standard Morse repulsion, the exponential repulsion does not become singular as *r* → 0, which eliminates potential instabilities in particle systems.

**Figure 13.**
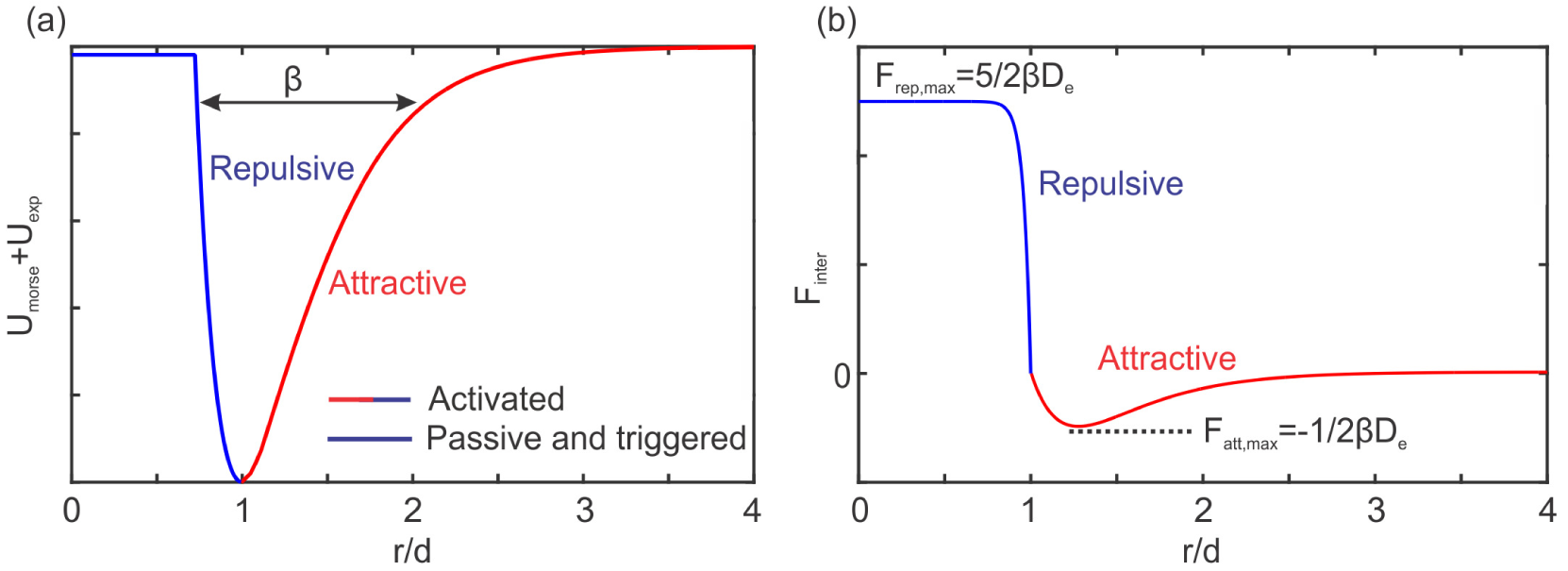
Schematic of the Morse+exponential potential (left) and the resulting adhesive force (right) used to mimic inter-platelet attractive/repulsive forces. Passive and triggered platelets only generate repulsive forces to prevent overlap, whereas activated platelets attract each other as well.

The value of β*d* defines the range of interactions between platelets; it is selected to be 2.5 to prevent overlap of passive platelets though partial overlap is allowed for activated platelets (*F_rep,max_/|F_att,max_*| = 5). The Morse attractive force is negligible for *r/d* > 3, which is considered as the cutoff radius for the attractive forces to avoid long-range interactions. The undetermined parameter *D_e_*, which controls the magnitude of platelet interaction forces, and *r_eff_*, which controls the number of particles in the simulation, are estimated by comparing the simulated thrombus size and shape against experimental observations^16^ under the same hemodynamic conditions. To calibrate the platelet aggregation model, we consider an interaction distance of 2*d* between platelets within which resting platelets may become activated.

#### Platelets and fibrinogen transport

We solve the transport of platelets and fibrinogen (denoted Fbg) using an advection/diffusion equation:

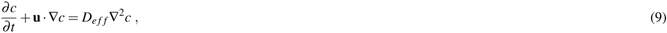

where *c* is the concentration field and *D_eff_* is the shear-induced effective diffusion coefficient. Note that the physiologic concentration of Fbg in blood is 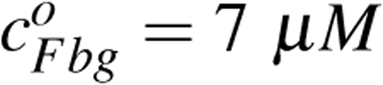^30^. Transport of platelets is modeled by treating platelet concentration as a continuum field, with ≈ 4 × 10^5^ platelets per *mm*^3^ of blood. The effective diffusion coefficient for platelets and macromolecules (*e.g*., Fbg) due to presence of red blood cells is taken as a function of the local shear rate based on an equation proposed by Wootton *et al*.^31^, 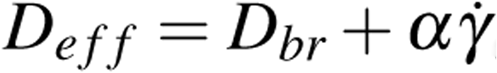, where the Brownian diffusion *D_br_* is 1.58 × 10^−13^ and 3.1 × 10^−11^ m^2^ /*s* for Fbg and platelets, and the coefficient α is 6.0 × 10^−14^ and 7.0 × 10^−13^ *m*^2^ for Fbg and platelets, respectively^31, 32^. In this work, we introduced two assumptions to reduce the complexity of the transport problem: first, we assume the value of shear rate inside the false lumen to be 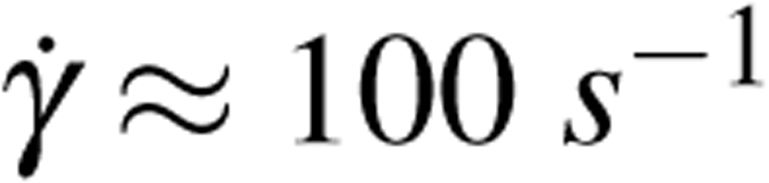 (see comments below in Discussion) to be consistent with flow simulation results herein and leads to a uniform value of diffusion coefficient; second, to avoid spurious solutions at high Peclet numbers (where *Pe* =*D_eff_ U_avg_*/v ≈ 10^6^, with the mean velocity *U_avg_* at inlet, and the blood kinematic viscosity ν), which requires special treatment in the numerical scheme and typically leads to long and expensive simulations to reach the desired static solutions, we increase the diffusion coefficients by two orders of magnitude. Hence, the final values of *D_eff_* are 6.16 × 10^−10^ and 1.01 × 10^−8^ *m*^2^/*s* for fibrinogen and platelets, respectively. All concentrations are normalized by physiologic values, with the initial distribution of Fbg and platelets assumed homogeneous everywhere.

#### Clot volume fraction estimation

To analyze the evolution in thrombus size and shape and to anticipate future simulations of multiphasic, long-term remodeling, we used a continuum representation of the adhered pseudo-platelets in the clot. This can be achieved easily in the context of the FCM by estimating the local volume fraction of FCM particles as:

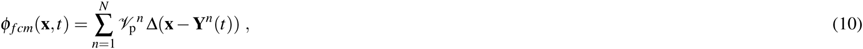

where 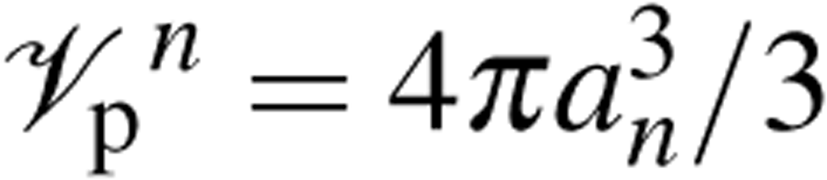 is the volume of each FCM particle. A larger standard deviation (σ ≈ 4*d*) in the Gaussian kernel in Eq. (4) yields a smooth representation of φ_*fcm*_.

Knowing the local concentration of fibrinogen and platelets, and using the estimated φ_*fcm*_ field, we can evaluate clot volume fraction φ*c* as:

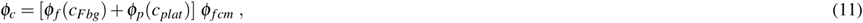

where an empirical correlation relates fibrin volume fraction φ_*f*_ to the concentration of Fbg, and φ_*p*_ denotes the platelet volume fraction^25^. Further details regarding the estimation of φ_*f*_ and φ_*p*_ can be found in the Discussion.

#### Flow and concentration boundary conditions

Mouse-specific, inlet flow rate waveforms were prescribed individually at the level of the proximal suprarenal aorta as a Dirichlet boundary condition (using a Womersley velocity profile^33^) based on pulsed wave Doppler measurements taken at the same location^16^. Two-parameter (also known as RC) Windkessel models were used to match PW Doppler measurements in the major outlet vessels (see Figure 14). A detailed study on robustness and stability of different outflow boundary conditions showed that two-parameter (RC) models are more stable than three-parameter (RCR) models^34^. The choice of two-parameter model implies less uncertainty as parameter estimation is not needed for the additional resistance.

**Figure 14.**
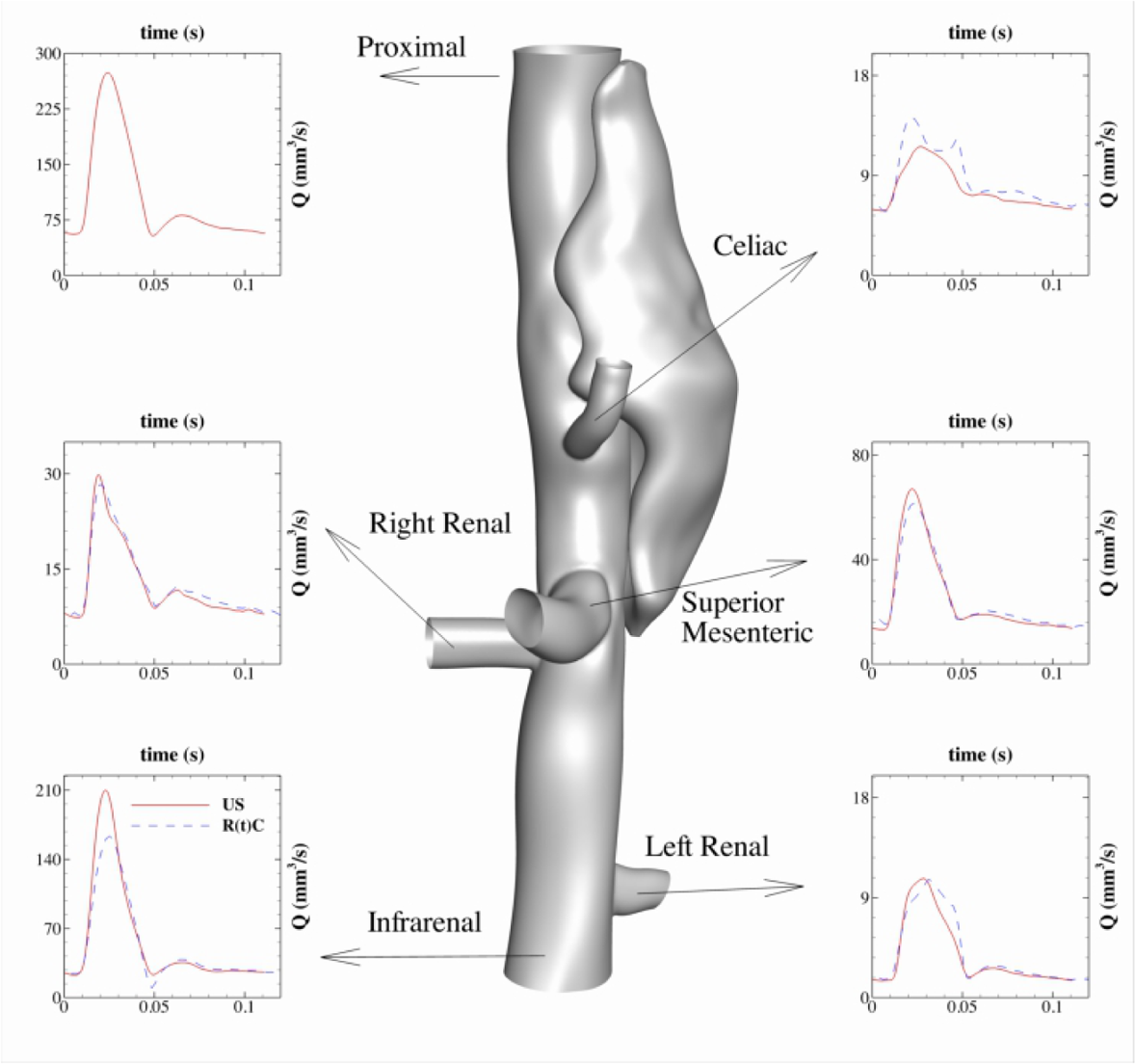
Inlet/outlet boundary conditions for the lesion with the medium false lumen: a Dirichlet flow waveform measured by ultrasound^16^ is imposed at the proximal inlet, while Windkessel RC type BCs are prescribed at every main outlet branch. A time-dependent resistance R(t) is estimated based on the experimental flow waveforms to improve the accuracy (shown in the inset plot for each outlet).

Using the average flow rates at each outlet and the flow analysis in a network of vessels given in^34^, we are able to calculate the steady RC values for each outlet. Our numerical observations, however, showed that the steady RC values do not perform well in the vasculature with the dissection, where the flow dynamics is significantly affected by the presence of the false lumen. Taking advantage of flow waveforms at the outlets, we computed time-dependent R(t)C values to reproduce the flow waveforms more effectively (shown in Figure 14). Further, at each outlet, a zero-flux Neumann boundary condition is considered for velocity and concentrations.

#### Data availability

All data generated or analyzed during this study are included in this published article (and its Supplementary Information files).

## Acknowledgements

This work was supported by NIH grant U01HL116323. Computational resources were provided by the Center for Computation and Visualization (CCV) at Brown University, TACC/STAMPEDE and SDSC/COMET through the XSEDE grant (TGDMS140007), and ANL and ORNL supercomputers through a DOE/INCITE award. The original computational domains and inlet/outlet conditions came from^16^, with particular thanks to Professor Craig Goergen at Purdue University.

## Author contributions statement

H.L. carried out the simulations and performed analysis; A.Y. performed analysis and participated in the design of the study, drafted the manuscript and the subsequent revisions; P.D.A performed image analysis and segmentation, and collaborated on the parameter estimation. M.R.B carried out the experiments and image segmentation; J.I. collaborated on the visualizations and created animations; J.D.H. helped design the study and helped draft and finalize the manuscript; G.E.K. helped design and coordinate the study and helped draft the manuscript. All authors gave final approval for publication.

## Additional information

The authors have no competing interests.

